# Active fluctuations of cytoplasmic actomyosin networks facilitate dynein-driven intracellular transport along microtubules

**DOI:** 10.1101/2024.05.23.595269

**Authors:** Takayuki Torisawa, Kei Saito, Ken’ya Furuta, Akatsuki Kimura

## Abstract

Inside cells, molecular motors transport cargoes within the actively fluctuating environment known as the cytoplasm. How fluctuations in the cytoplasm affect motor-driven transport is not fully understood. In this study, we investigated the role of fluctuations for transport along microtubules using *C. elegans* early embryos, focusing on transport driven by cytoplasmic dynein. An artificial motor-cargo complex showed faster transport *in vivo* than *in vitro*, suggesting an *in vivo* acceleration mechanism. We also found that endogenous early endosome transport by dynein is significantly enhanced by the fluctuations in the cytoplasm, which is attributed to the activity of actomyosin networks. An *in vitro* force measurement of dynein suggests that the asymmetric force response would play a key role in the acceleration. This study provides insights into a regulatory mechanism of molecular motors within actively fluctuating cytoplasm, potentially utilizing random force originating from fluctuating dynamics in the cytoplasm to increase transport efficiency.

## Introduction

Intracellular transport mediated by molecular motors is essential for various cellular processes, including neuronal development, organelle positioning, and cellular homeostasis. Microtubule-based motors, such as dynein and kinesins^1,2^, are involved in the transport and positioning of many types of cargos, such as endosomal organelles^3–5^, mitochondria^6^, and Golgi^7^. Dynein and kinesin also contribute to neuronal development through nuclear migration, and their dysfunctions cause neuronal diseases such as lissencephaly^8,9^. In addition, motor-driven intracellular transport not only moves organelle directly but also contributes to the nuclei positioning through mutual reaction force arising from vesicle transport in cytoplasm^10,11^.

Decades of *in vitro* and *in vivo* studies have brought deep insights into the motors’ walking and regulatory mechanisms. However, there remains a puzzling problem about the difference in the speed of motors between *in vitro* and *in vivo*: the speed from *in vitro* studies is mostly slower than the speed obtained from *in vivo* studies^12^. This difference is not apparent because most motor-driven transport occurs in the cytoplasm, an environment crowded with macromolecules^13–15^. It is known that crowding macromolecules make the cytoplasm highly viscous^16,17^. Cargos transported in a highly viscous environment experience greater drag force compared to those in typical buffer solutions used in *in vitro* studies. Given that a greater drag force generally slows down the moving object, it is reasonable to infer that the transport *in vivo* would be slower than *in vitro*, assuming that the maximum driving force remains constant. However, as mentioned above, previous *in vitro* and *in vivo* studies have suggested that motor-driven transport along microtubules is faster than *in vitro* in many cases^12,18,19^. Microtubule-based motors *in vitro* only achieve velocities up to 1 µm/s, whereas, in the case of early endosome transport in cells, velocities of several microns per second have been observed^20,21^, suggesting an acceleration mechanism within cellular environments. However, the previous observations are a mixture of *in vitro* and *in vivo* studies with various species and cell types, making it difficult to directly compare the motility in both situations.

Several possibilities have been considered to explain this apparent discrepancy in transport speed. One possible important factor is the difference in the number of motors involved in transport between *in vitro* and *in vivo*^22–24^. However, in previous *in vitro* experiments, where kinesin or dynein was aligned on DNA-origami nanotubes or double-strand DNA, the acceleration effect of increasing the motor number was limited^25–28^. Another possibility is that cellular modulations of the motor’s motility by regulatory proteins induce *in vivo* acceleration. In the case of cytoplasmic dynein, a minus-end directed microtubule-based motor, it is known that the regulatory proteins, such as LIS-1 and NDE1/NDEL1, modulate dynein’s motility^29,30^. It is also known that dynein-dynactin-adaptor complexes formed via adaptor proteins, such as Bicaudal D (BicD), Hook, and Spindly, promote unidirectional movement^2,27,31^. However, the studies observing these movements have not reported speeds comparable to *in vivo* dynein-driven transport.

Another possible explanation for the apparent discrepancy in transport speed is the contribution of a dynamic aspect of cytoplasmic environments: the cytoplasm is characterized not only as a crowded environment packed with high concentrations of macromolecules but also characterized by the existence of various active, energy-consuming processes. Such processes, including metabolism^32^ and fluctuating dynamics of cytoskeletal networks^33,34^, have been reported to produce forces in the cytoplasm and affect the physical properties of the intracellular environment^33,35^. Since all the active processes share the cytoplasm as their working space, the forces generated by one process can act as an environmental force for the molecules involved in other processes. Previous *in vitro* studies using the optical tweezer have revealed that external force can change the dissociation kinetics of microtubule-based motors^36–39^. Thus, the environmental forces from other processes would affect the motors’ activity. In addition, a recent study has also shown that the artificially generated non-Gaussian fluctuation, expected to mimic the actively fluctuating force in cells, accelerates the motor’s movements^40^. However, although previous studies have suggested that environmental forces can regulate motor-driven transport, the effects of the actively fluctuating forces on *in vivo* transport remain elusive.

This study aimed to reveal the role of fluctuating dynamics in the cytoplasm on motor-driven transport *in vivo*, focusing on the effects of actomyosin dynamics in the cytoplasm. Not only do actin and myosin networks produce contractile force in the cell cortex to induce the morphological dynamics of cells^41,42^, but they also form networks in the cytoplasm^43–46^. These cytoplasmic actin or actomyosin networks not only facilitate directed motion, such as the transport of the cell nucleus and chromosomes in mouse embryos^41–45^ and starfish oocytes^47,48^, but also generate active random forces within the cytoplasm, consequently influencing the physical properties of the cytoplasm^33^.

In this study, we utilized dynein-driven transport in *C. elegans* early embryos as our model system. We found that the dynein-cargo complex moved more rapidly *in vivo* than when observed *in vitro*, by directly comparing the motility of an artificial motor-cargo complex, which contains a minimum truncated construct of dynein, between *in vitro* and *in vivo.* This result suggests that the faster transport *in vivo* is not induced by the regulatory proteins in a cell, but by other cellular factors. By modulating the fluctuating dynamics of actomyosin networks within the cytoplasm, we found that actomyosin dynamics positively regulates the speed of endogenous dynein-driven transport in *C. elegans* embryos. Finally, combined with the asymmetric force response of dynein measured *in vitro*, we propose that the active fluctuations of the cytoplasmic actomyosin contribute to faster microtubule-based transport by dynein.

## Results

### Dynein-driven transport in living embryos is faster than *in vitro* situations

We constructed an experimental system where the motility of the same motor-cargo complex can be compared *in vitro* and *in vivo* (Figure 1A). To reduce the potential effects of endogenous *C. elegans* regulatory factors existing *in vivo*, we chose the truncated human dynein (hsDHC-1_382k_) (Figure S1A), which is a minimum motile construct including only the motor head and the linker domain^25,27,28,49^, as a transporting motor. Since most of the regulatory proteins of dynein bind to the tail region, by using the truncated tail-less dynein, we reduced the heterogeneity of regulatory states of dynein originating from the association with other proteins in *C. elegans* embryos. The SNAP-tagged hsDHC-1_382k_ were conjugated with the benzylguanine-modified 100-nm-beads (BG-beads). The size of the beads falls within the range of typical vesicles^50,51^. The artificial motor-cargo complex was introduced into *C. elegans* early embryos by injecting them into worm gonads^52^. We found the successful incorporation of the complexes into the embryos and their unidirectional movements (Figure 1B). The average transport speed of incorporated complexes was 6.9 ± 4.5 × 10^2^ nm/s (mean ± SD).

**Figure 1.**
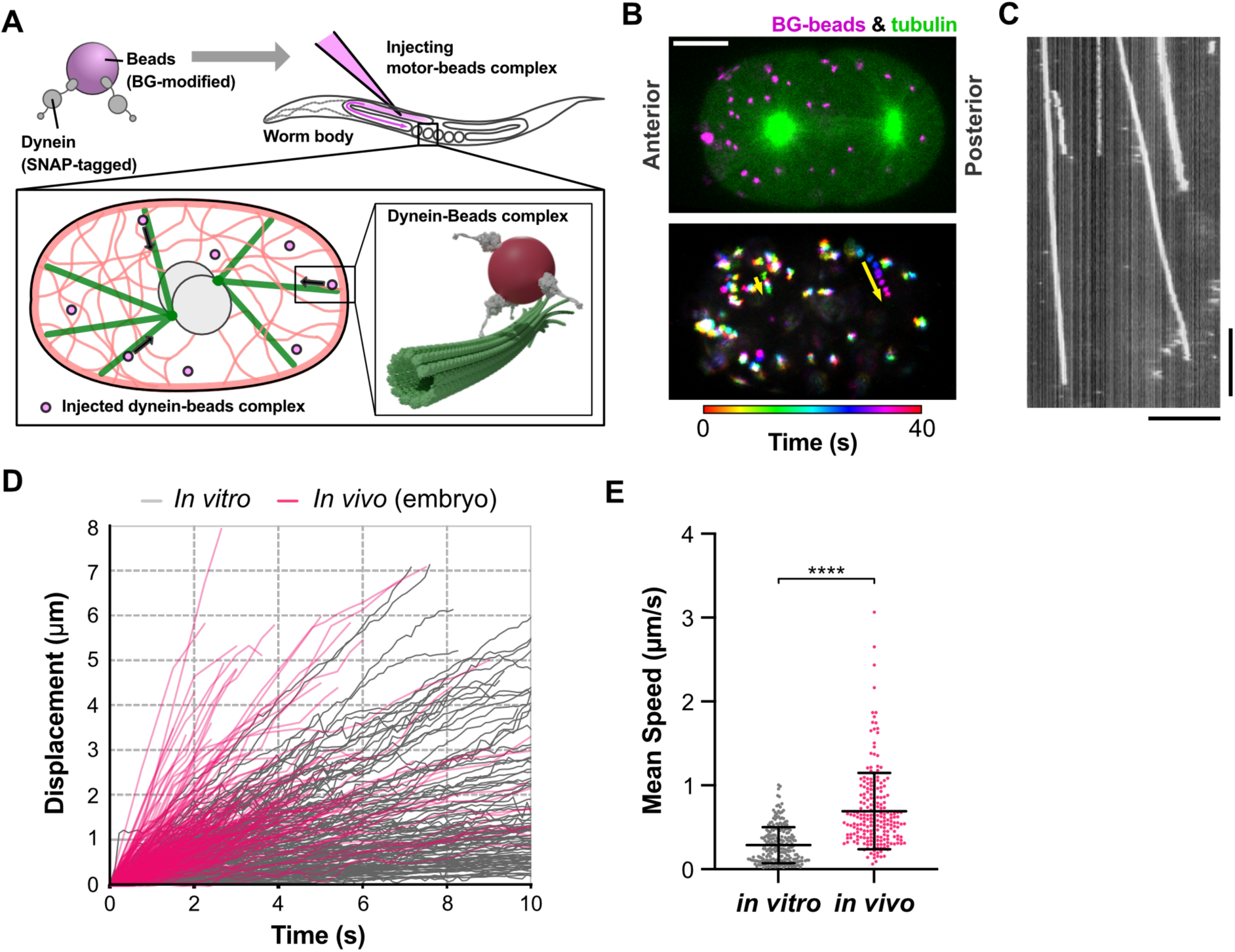
Direct motility comparison of artificial motor-cargo complex between *in vitro* and *in vivo*. (A) Schematic diagram of experiments for directly comparing the motility of the artificial motor-cargo complex between *in vitro* and *in vivo*. The SNAP-tagged minimum construct of the human dynein motor (1,306-4,646 aa) was conjugated covalently to the BG-modified fluorescent beads. The motility of motor-cargo complexes was analyzed *in vitro* using the typical experimental setup and *in vivo* by incorporating them into the embryo through the gonad injection. (B) The image of GFP-fused tubulin (green) and injected motor-beads complex (magenta) in a *C. elegans* embryo (left) and the temporarily-color-coded image showing the movements of the incorporated motor-beads complex. The yellow arrows indicate the unidirectional movements toward the centrosomes (not visualized). (C) Kymograph showing the movements of motor-cargo complexes along a taxol-stabilized rhodamine-labeled microtubule. Bars indicate 10 s and 10 µm, respectively. (D) Trajectories of individual motor-cargo complexes are shown. The trajectories are projected to the one-dimensional straight lines, which are obtained by the least square fitting to two-dimensional trajectories. The number of trajectories is 228 for *in vitro* movements (from 24 repeated experiments) and 215 for *in vivo* movements (from 20 embryos). The magenta and gray lines indicate the trajectories from *in vivo* and *in vitro* observations. (E) Comparison of mean speeds of the unidirectional movements of artificial motor-cargo complexes *in vitro* and *in vivo*. Individual plots show the mean speed of each trajectory, calculated by the linear fitting. Bars indicate Mean ± SD. The difference in mean transport speed was tested with Welch’s *t*-test (*p* < 0.0001).

We then examined the *in vitro* motility of the same samples used for *in vivo* observations. In a typical buffer environment (see Materials and Methods), we found that the average speed of dynein-bead complexes was 2.9 ± 2.1 × 10^2^ nm/s (mean ± SD), significantly slower than *in vivo* and comparable to those from previous *in vitro* studies^25,27,28^ (Figures 1C-1E). To exclude the possibility that the non-specifically attached endogenous motors drove the unidirectional movements of dynein-bead complexes in embryos, we observed the movements of beads without the conjugation with dynein. We found that the non-dynein-conjugated beads in the embryo did not exhibit unidirectional motion (Figures S1C and S1D), indicating that the endogenous dynein did not involve the unidirectional movements of the hsDHC-1_382k_-conjugated BG-beads. We also found that the movements of BG-beads with dynein were less diffusive than that without dynein (Figure S1E). This difference in diffusivity is supposed to be caused by the specific interaction of bead-dynein complexes with microtubules mediated by the conjugated dynein. Collectively, our results demonstrate that the motor-cargo complex is able to move faster *in vivo* than *in vitro*, even when we use the same sample in both situations.

Our direct comparison of motility showed that dynein-driven transport in the cytoplasmic environment was faster than in a typical *in vitro* condition. This result clearly demonstrates that transport *in vivo* is faster than *in vitro*, as suggested by previous studies. Note that our experiments used the truncated dynein from other species, which is expected not to be affected by endogenous regulatory proteins, suggesting the importance of other cellular factors for faster transport *in vivo*. We then hypothesized that the active fluctuations in the cytoplasm, originating from the motion and force produced by the energy-consuming cellular processes, facilitated the transport. Among the fluctuation-producing cellular processes, we decided to focus on the fluctuating dynamics of actomyosin networks in the cytoplasm because they are present throughout the cytoplasm and known to produce spatiotemporally random forces^33^.

### Early endosome transport in *C. elegans* embryos as a model system of *in vivo* transport exclusively driven by cytoplasmic dynein

To investigate the effects of fluctuating dynamics of actomyosin networks on intracellular transport along microtubules, we analyzed the transport *in vivo* under the perturbations on actomyosin. For this purpose, we focused on the intracellular transport of early endosomes, known as one of the main cargoes of cytoplasmic dynein^2,3^, and used the 1-cell stage *C. elegans* embryos as a model system (Figures 2A and 2B). We found that early endosomes moved at 8.1 ± 3.8×10^2^ nm/s (mean ± SD; Figures 2C and 2D), comparable to the transport in the previous studies^20,21^. In our observations, all the transports of early endosomes were towards the embryo center, suggesting that the transport was mediated mainly by dynein. Furthermore, we found almost no unidirectional early endosome transport in the dynein-depleted embryo where the endogenous dynein was degraded by the auxin-induced degron (AID) system (Figures 2E-2G), which degrades the degron-tagged protein specifically upon the drug treatment^53–55^. These results show that the early endosome transport in *C. elegans* embryos, at least in 1-cell stage embryos, is exclusively driven by dynein, in contrast to other systems where a minus-end directed kinesin contributes to the transport^56^.

**Figure 2.**
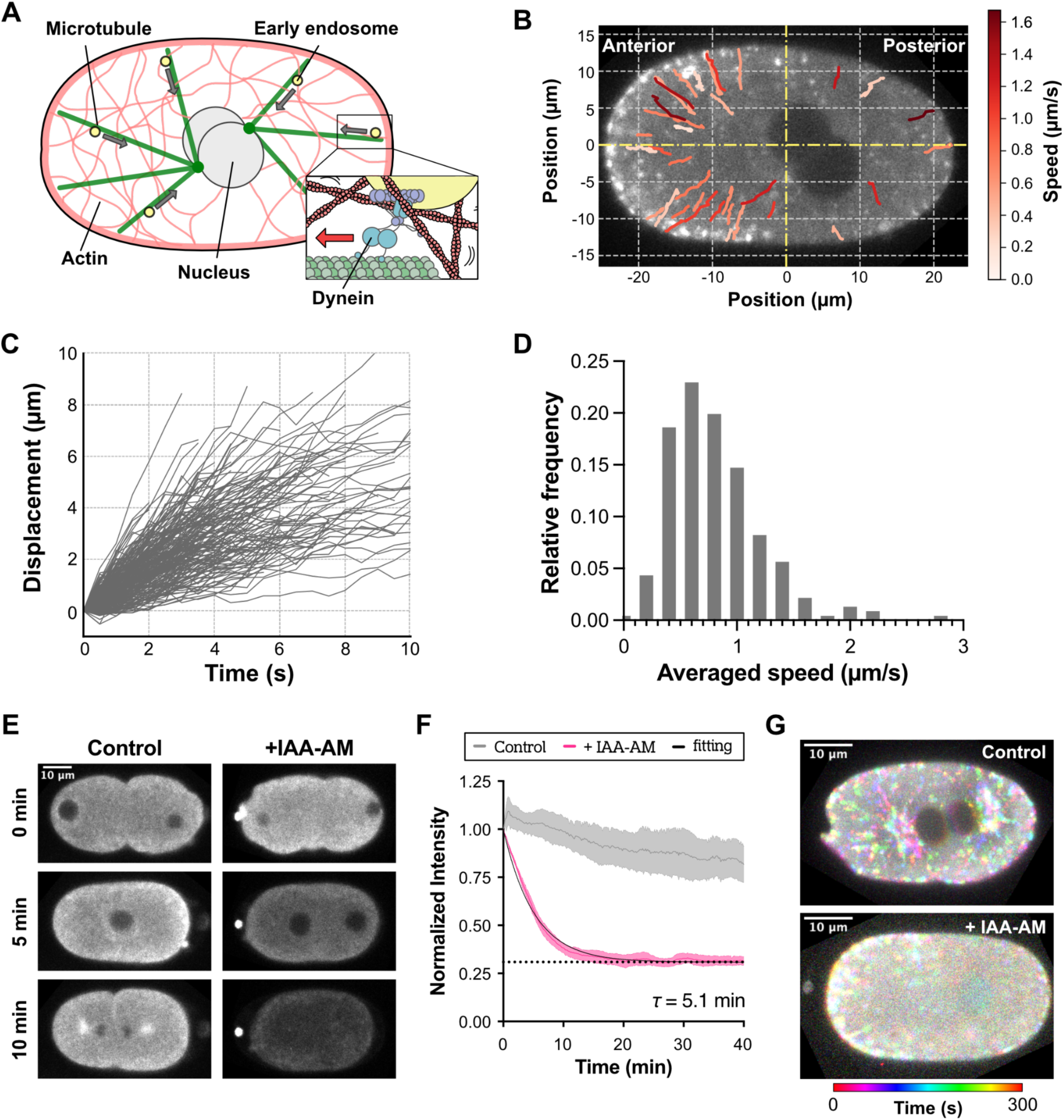
Early endosome transport in *C. elegans* 1-cell stage embryo is driven by cytoplasmic dynein. (A) Schematic diagram of early endosome transport in *C. elegans* embryo. Cytoplasmic dynein transports early endosome vesicles from the peripheral region of the embryo to the centrosomes attached to the pronuclei. (B) A typical image of a 1-cell stage embryo obtained from the strain exogenously expressing RAB-5::GFP. The lines indicate the two-dimensional trajectories of individual RAB-5 positive early endosomes. The mean speed of each trajectory is represented by the colors as indicated by the bar. (C) Trajectories of early endosomes projected to the one-dimensional straight lines. The number of trajectories is 348, obtained from 13 embryos. (D) The distribution of mean transport speed. The speeds were calculated from individual trajectories, which are shown in (C). (E) Time-lapse images of 1-cell stage embryos endogenously expressing the degron-fused dynein heavy chain (DHC-1::degron::GFP) with/without 5 mM IAA-AM. The time elapsed after starting the observations is indicated. (F) Time series of the intensity of degron-fused DHC-1 in the control (gray) and the +IAA-AM (magenta) conditions. The black line represents the single-phase exponential decay fitted to the +IAA-AM condition. The characteristic decay time (τ) is shown in the plot. (G) Temporally-colored images of 1-cell stage embryos expressing RAB-5::mCherry and the degron-fused DHC-1 with/without 5 mM IAA-AM.

Before perturbing the actomyosin dynamics, we examined the relationship between dynein-driven transport and the actomyosin dynamics. During the observations, we noticed that some early endosomes showed correlated motion with the deformation of actin networks (Figures S2A and S2B), indicating the direct interaction of early endosomes and actomyosin networks^21^. Based on this result, we further analyzed the relationship between early endosome transport and actomyosin networks. We found a coincidence between the a symmetry of transport speed and the asymmetry of actomyosin fluctuations: the transport speed of early endosomes on the posterior side was significantly higher than that on the anterior side (Figure S2C), and the fluctuation of actin networks, which was quantified by the optical flow analysis, was more prominent on the posterior side (Figure S2D). However, at the same time, we found an asymmetry in actin density along the anterior-posterior axis (Figure S2E and S2F). Because the density of actomyosin networks affects the transport speed by changing the drag force to the cargo-motor complex, it was difficult to determine whether the fluctuation or density of actin networks is more critical for dynein-driven transport, motivating the analysis of the transport under the perturbations on the actomyosin activity.

### Speed of dynein-driven intracellular transport depends on the degree of fluctuating dynamics of actomyosin networks in the cytoplasm

To perturb the activity of actomyosin networks, we adapted a genetic downregulation and a biochemical upregulation of *nmy-2*, the *C. elegans* non-muscle myosin II^57^, that generates force in the actomyosin network (Figure 3A). For the downregulation, we reduced the amount of NMY-2 in the embryo by RNAi, expecting it to weaken forces causing fluctuating dynamics of actomyosin networks. For the upregulation, we increased the embryonic NMY-2 level by injecting the purified NMY-2 complex through the gonad injection^52^.

**Figure 3.**
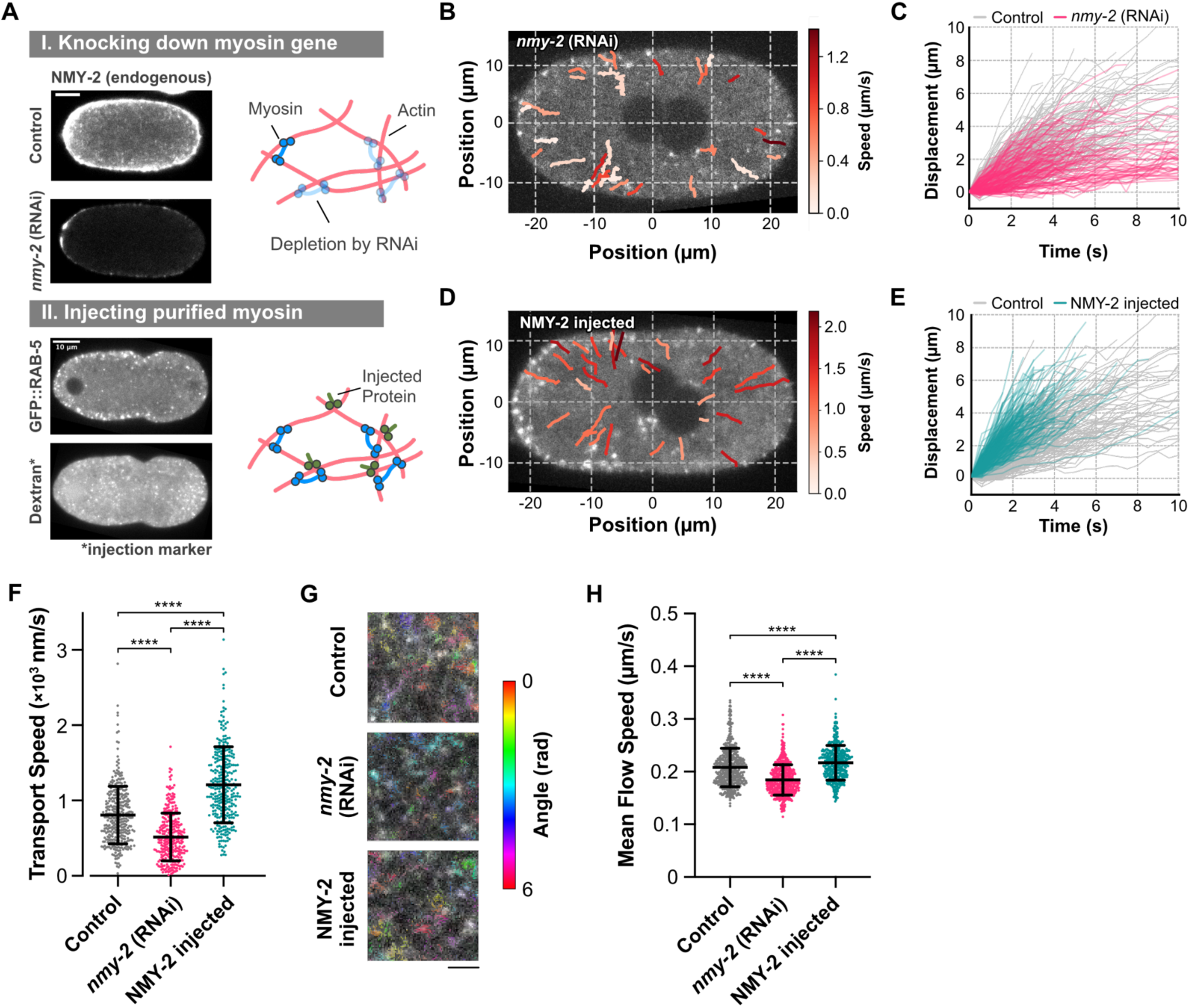
Speed of dynein-driven intracellular transport depends on the degree of fluctuating dynamics of actomyosin networks. (A) Schematic diagram showing the methods of up-and down-regulation of NMY-2 in the embryos. (top) For the down-regulation, the expression level of *nmy-2* was reduced by RNAi. The images show the typical example of the endogenously-tagged GFP::NMY-2 in the *C. elegans* early embryos from the control and *nmy-2* (RNAi) worms (top left). The ranges of brightness of the images are the same for both images. Knocking down *nmy-2* gene is expected to reduce the number of force generators in the actomyosin network (top right). (bottom) For the up-regulation, the amount of NMY-2 was increased by incorporating the purified proteins through the gonad injection. The images show the typical example of GFP::RAB-5 and the Texas Red-labeled dextran co-injected with NMY-2 complexes as an injection marker in the same embryo (bottom left). Injecting NMY-2 molecules is expected to increase the number of force generators in the actomyosin network (bottom right). (B) Typical image of a 1-cell stage *nmy-2* (RNAi) embryo obtained from the worm exogenously expressing GFP::RAB-5. The lines indicate the two-dimensional trajectories of individual RAB-5 positive vesicles that moved unidirectionally during 5-minute observations. The mean speed of each trajectory is represented by the colors as indicated by the bar. (C) Comparison of the one-dimensionally projected trajectories of RAB-5 positive vesicles in the control (gray) and the *nmy-2* (RNAi) (magenta) embryos. The data of the control embryos is the same as shown in Figure 1. The number of trajectories for the *nmy-2* (RNAi) embryos is 315, obtained from 14 embryos. (D) Typical image of a 1-cell stage NMY-2 injected embryo with the two-dimensional trajectories of unidirectionally moving RAB-5 positive vesicles. The line colors are the mean transport speed indicated by the color bar. (E) Comparison of the one-dimensionally projected trajectories of RAB-5 positive vesicles in the NMY-2 injected embryos (cyan). The number of trajectories from the NMY-2 injected condition was 311, obtained from 11 embryos. (F) Comparison of mean transport speed of RAB-5 positive vesicles in the unperturbed (control), *nmy-2* (RNAi), and NMY-2 injected embryos. Each plot represents the mean speed from individual trajectories. The bars indicate mean ± SD. (G) Flow fields imposed on the actin images from the unperturbed (control), *nmy-2* (RNAi), and NMY-2 injected embryos. The color indicates the flow vectors’ angle, as shown in the right color bar. The scale bar indicates 10 µm. (H) Comparison of mean flow speed of actin networks obtained from the optical flow analysis. Each plot represents the mean flow speed calculated from the individual pairs of images. The bars indicate the mean and SD. The number of embryos analyzed was 10 for the unperturbed (control), 10 for *nmy-2* (RNAi), and 8 for NMY-2 injected conditions, respectively.

We first analyzed the motile property of early endosome transport under the downregulated condition of NMY-2. In the *nmy-2* (RNAi) embryos, we found that our RNAi condition reduced the amount of NMY-2 to about 24% (Figures S3A) and that the transport speed of early endosomes decreased to less than 70% (5.2 ± 3.2 ×10^2^ nm/s; Figures 3B, 3C, and 3F). We also found that the presence of a myosin II inhibitor, blebbistatin, resulted in a speed reduction (Figure S3B). These results suggest that the activity of actomyosin regulates the activity of dynein-driven transport.

We then analyzed the motile property of early endosomes under the upregulated condition of NMY-2. We incorporated the purified NMY-2 complex into the embryo by injecting the protein molecules into the worm gonad^58^. We purified the *C. elegans* NMY-2 complex containing the HMM fragment, the full-length light chain 4 (MLC-4), and the full-length light chain 5 (MLC-5), following the previous study^59^ (Figure S3C). In the NMY-2 injected embryos, we found that the transport speed of early endosomes was faster than that in the unperturbed embryos (12.1 ± 5.0 ×10^2^ nm/s, mean ± SD; Figures 3D-3F), in contrast to the NMY-2 depleted embryos. These results suggest a correlation between the actomyosin dynamics and the speed of dynein-driven transport. However, we were concerned about the possibility that the perturbations on NMY-2 changed the size of early endosomes^60^, which affects the drag force on the cargo-motor complex and the transport speed. Assuming the amount of RAB-5 is proportional to the size of early endosomes, we analyzed the early endosome size by measuring the intensity of GFP::RAB-5 (Figure S3D). The distributions of RAB-5 intensity showed a significant difference among the unperturbed and NMY-2 perturbed embryos (Figure S3E). However, when we compared the transport speeds of early endosomes with similar intensity values, we found that the transport was faster in NMY-2 injected embryos and slower in *nmy-2* (RNAi) embryos, regardless of the size of early endosomes (Figures S3F). These results suggest that the amount of NMY-2, not the size of early endosomes, positively affects the dynein-driven transport of early endosomes.

Since the amount of force-generating NMY-2 is expected to correlate with the fluctuating dynamics of actomyosin networks, we analyzed the degree of fluctuations of actomyosin networks by calculating the optical flow of actin signal in the cytoplasm (Figures 3G and S3G). We found that the mean flow of actin networks changed depending on the perturbations on NMY-2 (Figure 3H), indicating a positive correlation between the degree of actin fluctuation and the speed of dynein-driven transport. To exclude the possibility that the perturbations on NMY-2 caused a change in the mesh structure of actomyosin, which can affect the drag force on transport and its speed, we analyzed the effect of NMY-2 perturbations on the spatial pattern of actin in the cytoplasm visualized by GFP::utrophin^61^. We calculated the coefficient of variations (CV)^62^, the ratio of the standard deviation to the mean, of actin intensity and its dependency on the window size (Figure S3H). The dependency on the window size is expected to reflect the characteristic length of actin networks because if the window size is smaller than the characteristic size, the CV will change according to the window size, but if the window size becomes larger than the characteristic size, the CV will not vary largely. Note that if the pixel-shuffled image is analyzed, where the positional order of all pixels is randomized, the CV is expected to reach saturation rapidly because the randomization disrupts the characteristic structure of the network and decreases the characteristic length. We found that the CV of actin intensity increases with window size according to the characteristic network density, and the decrease and increase of NMY-2 level in the embryo did not induce any apparent changes in the window size dependency (Figure S3I). This result suggests that the perturbations on NMY-2 have little effect on the network structure of actin in the cytoplasm. To check our method, we analyzed the pixel-shuffled image and confirmed that the CV increases faster than the original image. Our results indicate that the degree of fluctuating dynamics of actomyosin is a critical regulatory factor in the activity of microtubule-based transport.

The above observations using a fluorescent probe attached to dynein-driven early endosomes indicate that the fluctuations of actomyosin networks are coupled with the speed of dynein-driven transport along microtubules, suggesting that the active random forces from the actomyosin facilitate the transport. To directly evaluate the effect of fluctuating dynamics of actomyosin networks on the movement of materials within the cytoplasm, we analyzed the movement of cargo-size particles incorporated into the *C. elegans* embryo.

### Fluctuations in the cytoplasm changed depending on the perturbations of myosin

To investigate how the fluctuating dynamics of actomyosin networks in the embryo evoke fluctuations in the cytoplasm, we analyzed how the movements of materials in the cytoplasm are affected by the actomyosin dynamics (Figures 4A and 4B). We used the PEG-coated fluorescently labeled bead with a diameter of 100 nm, which is expected not to associate cellular materials non-specifically, to examine how actomyosin dynamics affects the random motion in the cytoplasm. The beads were incorporated into the embryos through the gonad injection^52,63^, under the control and the NMY-2 perturbed conditions (Figure S4**A**). To examine the effect of modulating embryonic NMY-2 level, we calculated the mean squared displacement (MSD) of the movements of beads (Figure 4C). We found a clear difference in bead diffusivity, as shown in the slope of the MSD plots, depending on the embryonic NMY-2 level. In the *nmy-2* (RNAi) embryos, the diffusion coefficient (*D*) of beads decreased to approximately 53% of that of the control embryos (*D* = 5.0 ± 0.3 ×10^4^ nm^2^/s for control and 2.7 ± 0.2 ×10^4^ nm^2^/s for *nmy-2* (RNAi), mean ± SEM). Note that we confirmed that the analyzed trajectories reflected the movements of single beads, not including the aggregated ones, by confirming that the intensity distributions of beads have a single peak (Figure S4B). On the other hand, in the NMY-2 injected embryos, we found that the bead diffusivity increased by approximately 70% (*D* = 8.6 ± 0.6 ×10^4^ nm^2^/s). Since the fluctuating component in the motion originates from the environmental force, these results suggest that the fluctuating forces in the cytoplasm correlated with the embryonic NMY-2 level. In addition, we found that the MSD exponents from the log-log plots were not significantly different among the control and the perturbed conditions (Figure S4C). Since the MSD exponent is affected by environmental confinement, we speculated that the mesh structure of actomyosin networks did not change upon the perturbations on NMY-2, consistent with the CV analysis of actin images (Figure S3I). We then hypothesized that the difference in bead diffusivity reflected the forces exerted by fluctuating actomyosin networks surrounding the beads, which is also suggested by a non-Gaussian characteristic of bead displacements (Figure S4D). However, we could not exclude the possibility that the environmental viscosity originating from the mechanical properties of actomyosin meshes changed depending on the perturbations on NMY-2.

**Figure 4.**
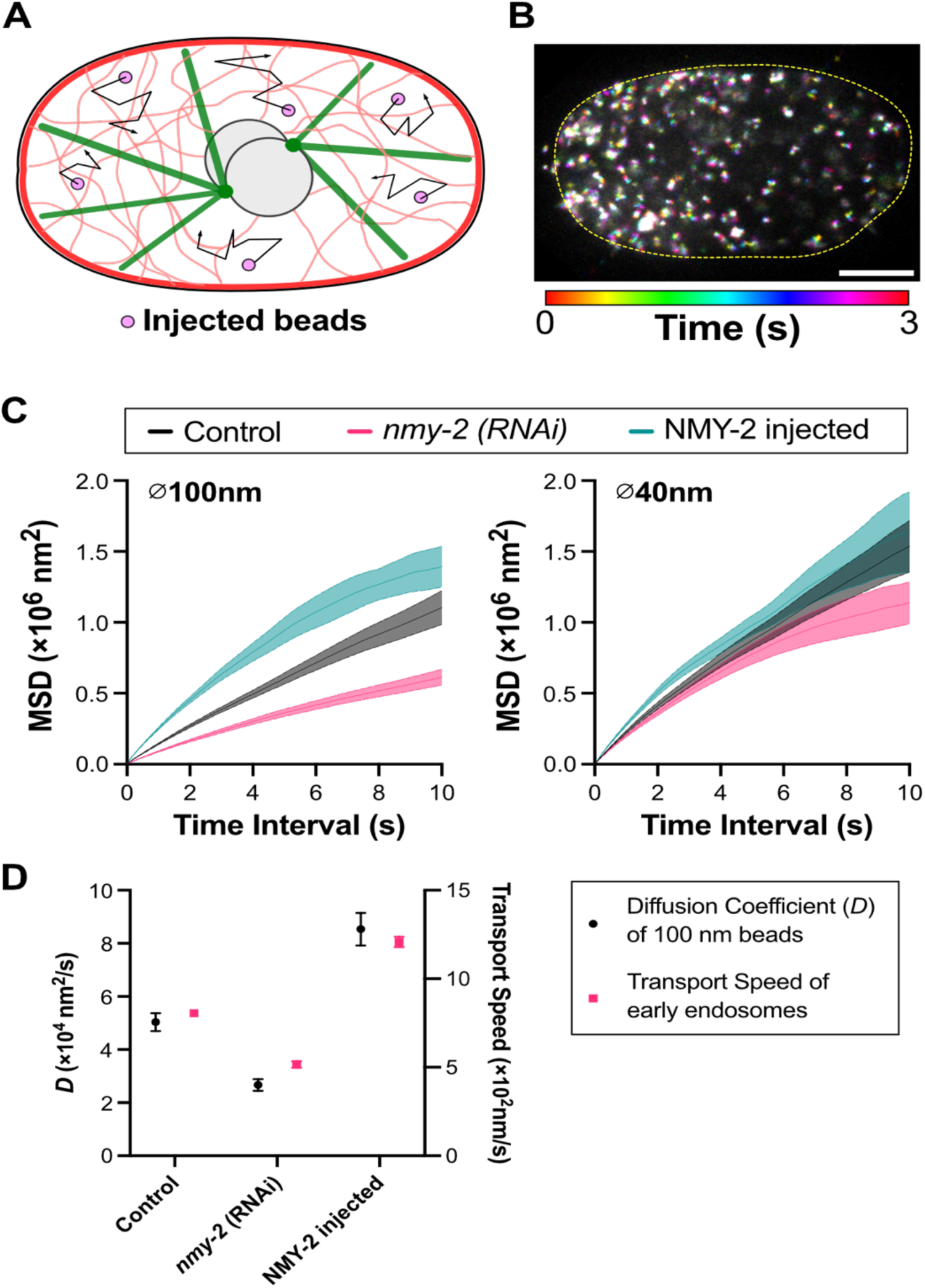
Effects of NMY-2 level in the *C. elegans* embryos on the movements of tracer particles. (A) Schematic diagram of the experiments, where fluorescent beads with 100 nm or 40 nm diameter are incorporated into the embryo through the gonad injection. (B) A temporal-color-coded image displaying the movements of the beads with a diameter of 100 nm in a 1-cell stage embryo. The time is indicated by the color, as shown in the bar below the image. (C) Averaged MSD plots of beads with a diameters of 100 nm (left) and 40 nm (right) obtained from the movements under the control (gray), *nmy-2* (RNAi) (red), and NMY-2 injected (cyan) conditions. Mean and SEM are shown. The number of trajectories used for the calculation was 339 from 10 embryos (100 nm, control), 346 from 13 embryos (100 nm, *nmy-2* (RNAi)), 331 from 10 embryos (100 nm, NMY-2 injected), 334 from 4 embryos (40 nm, control), 323 from 6 embryos (40 nm, *nmy-2* (RNAi)), and 387 from 7 embryos (40 nm, NMY-2 injected), respectively. (D) Comparison of diffusion coefficients of beads with the diameter of 100 nm (black) and the transport speed of early endosomes (red) under the unperturbed (control), *nmy-2* (RNAi), and NMY-2 injected embryos. Bars indicate SEM. The diffusion coefficients were calculated from the data shown in (C).

To address this point, we observed the movements of particles with a smaller size than the characteristic size of actomyosin meshes. Although there is no measurement of actomyosin mesh size in *C. elegans* embryos, several studies using other systems showed that the characteristic size was about 50 nm^64–66^. We then analyzed the movements of 40 nm beads, expecting that the size was below the mesh size. By calculating the MSD plot of 40 nm beads, we found that the differences in the diffusivity of beads among the conditions were not prominent in the case of 40 nm (Figure 4C; *D* = 8.7 ± 0.6 ×10^4^ nm^2^/s for control, 7.3 ± 0.5 ×10^4^ nm^2^/s for *nmy-2* (RNAi), and 10.1 ± 0.5 ×10^4^ nm^2^/s for NMY-2 injected). Based on these results, we thought that environmental viscosity was not the dominant factor, at least in the short time scale comparable to transport events (several seconds), but the active forces provided by fluctuating dynamics of actomyosin networks were critical for regulating transport. Furthermore, by comparing the dependency of the diffusivity of 100 nm diameter beads and the transport speeds of early endosomes on the NMY-2 level in embryos, we discovered a positive correlation between these two quantities (Figure 4D). This finding supports our hypothesis that the randomly fluctuating force within the cytoplasm, part of which originates from the activity of the actomyosin networks, influences the dynein-driven transport along microtubules.

### External force can induce an acceleration of dynein-driven transport through asymmetric force response of dynein

Our observations suggest that cellular environments accelerate dynein-driven transport along microtubules with randomly fluctuating forces in the cytoplasm. How is such an acceleration *in vivo* achieved? We hypothesized that how dynein responds to external force is a crucial factor. Several studies reported an asymmetric force response of cytoplasmic dynein I of *Saccharomyces cerevisiae*: dynein changes its dissociation rate from a microtubule depending on the direction of external forces^36,38^. This asymmetric force response of motors to environmental forces can be a potential mechanism of the acceleration of transport under the active fluctuations *in vivo.* Even if the direction of the external force is random, the asymmetric force response can induce a direction-dependent, asymmetric binding and unbinding from a microtubule, which may result in more frequent forward motion than without the external force. Based on this view, we investigated whether the human dynein used in our experiments showed a similar asymmetric force response. As demonstrated in the previous studies^36,38,39^, we conjugated hsDHC-1_382k_ to beads and exerted external load to dynein using the optical tweezer (Figures 5A and 5B). In the experiments, we moved the center of an optical trap back and forth along a microtubule. We note that when we pulled the trapped beads at a low loading rate, the strain to dynein would not reach sufficient strength before detachment, primarily because of the rotational freedom of the trapped bead. To avoid this, we pulled the beads at sufficiently high loading rates. We rapidly moved the beads in ∼70 µs and measured the duration of the binding events, occasionally observed during forward and backward movements of the trap center (Figures S5A and S5B). Like the yeast counterpart, we found that human dynein exhibits an asymmetric force response depending on the direction of applied forces. It easily detached from microtubules when pulled toward the minus-end, whereas it tended to bind strongly to microtubules when pulled toward the plus-end (Figures 5C and S5C). We also found that the frequency of binding events showed a significant difference depending on the direction of load (Figures S5C and S5D), where the binding of dynein to microtubule occurred more frequently during plus-end directed (backward) movements of the trap center than minus-end (forward) movements. Furthermore, the rate of strong binding events to all binding events depended on the loading rate and the direction of applied force: the faster movements of the trap center tended to induce more frequent strong binding of dynein (Figure S5E). The characteristic asymmetry in force response of dynein was pronounced with larger external forces, around approximately 6 pN, but was even observed in the range of weaker forces, around 1 pN, comparable to the active force measured in human cancer cells^67^. These results suggest that active fluctuating force in the cytoplasm can be harnessed by dynein’s asymmetric force response property and utilized for facilitating unidirectional transport in the cell (Figure 5D).

**Figure 5.**
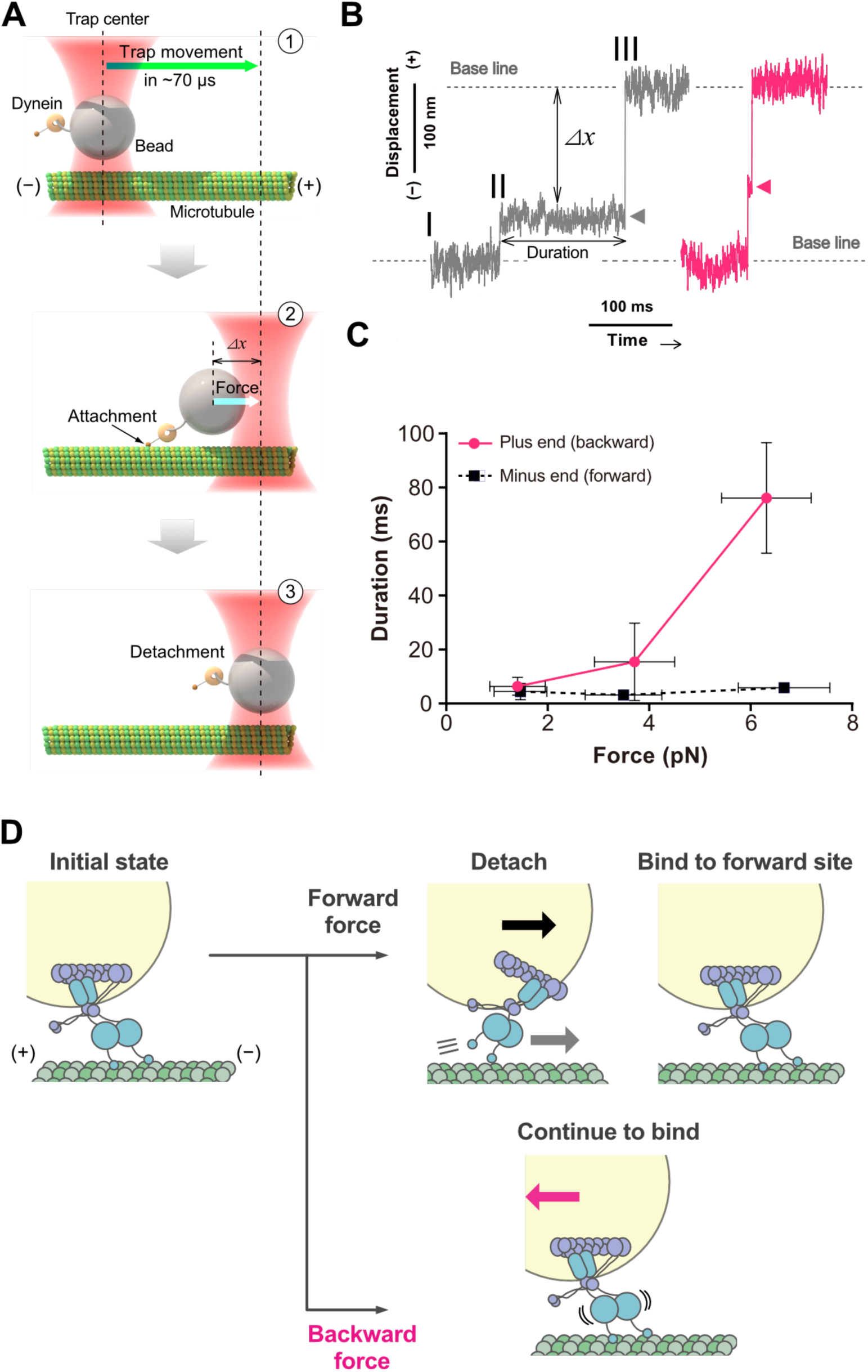
In vitro measurement of force response of optically-trapped dynein. (A) Schematic diagram of the experiment measuring the force response of dynein molecules. The signs denote the plus-end and minus-end of the microtubule. (B) Typical time series of a bead center. The center rapidly moves towards the plus-end of a microtubule (I to III). The duration of transient binding events of dynein during the scanning (II) was analyzed. The arrowheads indicate the long (purple) and short (blue) binding events. (C) Duration of strong binding events versus applied force. The data were acquired at a loading rate of 0.127 pN/µs (1 mM ATP). The duration data were classified into three groups according to the force (0–2.5 pN, 2.5–5.0 pN, >5.0 pN), and the sum of two exponential functions fitted the data in each group (n = 92–174). The longer time constants obtained from each fit were plotted against the mean force calculated in each group. The horizontal error bars represent the SDs of the raw data, and the vertical error bars denote the SEs, which were calculated as the SD of fitted parameters derived from bootstrapped data sets. The red solid line shows the duration when the plus-end-directed, backward load was applied to dynein. The blue dashed line shows the duration when the minus-end-directed forward load was applied. (D) Model of forward movement of dynein induced by external force and the asymmetric force response. The signs denote the plus-end and minus-end of the microtubule. If the external force is forward-directed, dynein is easily detached from a microtubule, stochastically rebind, and produces a forward step. The process contains a transition from the strong to weak binding state and is supposed to be coupled with the ATP hydrolysis cycle of dynein. If the external force is backward-directed, dynein tends to continue the strong binding state and does not produce a backward step. In cells, the force originating from other energy-consuming processes, such as actomyosin fluctuations, is supposed to act as an external force to dynein.

## Discussion

Intracellular transport of materials driven by molecular motors occurs within the cytoplasmic environment of the cell. Many force-generating processes co-occur within this environment, yet the impact of such environmental active processes on motor-driven transport still needs to be understood. We observed intracellular transport by cytoplasmic dynein in *C. elegans* early embryos. We found that fluctuations generated by actomyosin networks in the cytoplasm facilitate dynein-driven transport along microtubules.

Our observations revealed that the motility of an artificial motor-cargo complex was significantly enhanced in a cellular environment, indicating that transport *in vivo* is faster than *in vitro* (Figure 1E). In addition, we also found that the speed of dynein-driven early endosome transport changed as we increased or decreased the embryonic level of non-muscle myosin II (NMY-2) (Figures 3B-3F). By analyzing the spatiotemporal fluctuation of cytoplasmic actin networks, we found that the embryonic level of NMY-2 positively correlated with the active dynamics of actomyosin networks in the cytoplasm (Figure 3H), which was also correlated with the diffusivity of injected beads (Figures 4C and 4D). These results suggest that fluctuating dynamics of actomyosin networks led to the change in the randomly fluctuating force in the cytoplasm and the consequent change in the speed of dynein-driven transport.

Based on an *in vitro* force measurement of dynein, we hypothesized that *in vivo* acceleration of dynein-driven transport is mediated by the asymmetric force response of dynein (Figure 5C). In our model, environmental forces from the cytoplasmic actomyosin networks can be rectified by the cargo-attached dynein, leading to displacement following the force direction (Figure 5D). It is worth noting that a previous study has indicated that dynein harnesses active fluctuations of microtubules to achieve efficient transport^39^. This suggests that apart from the direct effects of actomyosin dynamics, there can be an indirect effect on dynein-driven transport by inducing microtubule fluctuations in the cytoplasm. In addition to previous studies on yeast dynein, our observations revealed that human dynein also exhibits an asymmetric force response. While differences in motility, including speed, affinity to microtubules, and diffusivity, between yeast and human dynein have been reported^25,27,28^, we found a qualitative agreement in their force responses, suggesting a possible universality of this asymmetric response of dynein. This possible universal force response of dynein may contribute to efficient transport in highly viscous cytoplasmic environments, typically expected to slow down transport, not only in *C. elegans* but also in other species.

The acceleration mechanism by rectifying external force may work strongly when the molecular motor forms a complex bound to cargo. Our experiments with injected beads revealed that the effects of NMY-2 perturbations varied with the size of the beads (Figure 4C). Beads of 40 nm diameter, comparable to the size of full-length dynein^68^, showed a weaker change in MSD plots depending on the perturbations on NMY-2. On the other hand, beads of 100 nm diameter, similar in size of smaller endosomes^50^, exhibited notable changes upon myosin II perturbations (Figure 4C). This contrast may be caused by the characteristic mesh size of the cytoplasmic actomyosin networks, previously reported to be around 50 nm in mammalian cells^64,69^. Although the mesh size of actomyosin networks in *C. elegans* early embryos has not been directly measured, our results suggest that the characteristic mesh size of *C. elegans* cytoplasmic actomyosin falls in a similar range. Future studies to capture fine structures of actomyosin networks in embryos will bring further insight into the characteristic mesh size and its role in transport regulation.

Previous studies, including ours, have shown that injecting materials into *C. elegans* gonads allow *in vivo* observations of material dynamics in embryos^52,63^, which can be compared with *in vitro* experiments. In this study, by injecting covalently-conjugated motor-cargo complexes, we observed faster movements *in vivo* than *in vitro* (Figure 1). Future studies using DNA nanostructures like DNA-origami will enable a more detailed examination of the artificial motor-cargo complexes^26,28,70^, including the impact of motor number and geometric configuration on cargo transport.

The acceleration of dynein-driven transport in a randomly fluctuating cytoplasm raises a question about its biological implications. One possibility, as mentioned earlier, is that dynein molecules are inherently equipped with a mechanism rectifying external force to efficiently transport cargo even in a high-viscosity environment, typically expected to slow down the transport of large vesicles^71,72^. Another perspective involves the energetics of the cell. The force generated by the ATP-consuming dynamics of actomyosin networks may be utilized not only to drive actomyosin dynamics itself but also to facilitate dynein-driven transport along microtubules through the asymmetric force response of dynein. In addition, this byproduct of randomly fluctuating dynamics of actomyosin could result in the unidirectional transport of larger cellular components. In *C. elegans* embryos, dynein-driven transport along a microtubule produces the mutual reaction force, called cytoplasmic pulling force, which pulls the nucleus through the centrosome^10,12,73–75^. Thus, the observed acceleration of microtubule-based transport, caused by randomly fluctuating dynamics of actomyosin, may contribute to the efficient transport of the nucleus, even in a cellular context like *C. elegans* embryos, where the direct contribution of actomyosin is not apparent. Our results suggest a mechanism where the inevitable byproduct of cytoplasmic fluctuations is converted into unidirectional motion of small vesicles and large organelle. We expect that future comprehensive studies, including live cell observations in other species, *in vitro* experiments, and *in vivo* observations of elements that can be characterized *in vitro*, will likely elucidate how the active cytoplasmic environment impacts the cellular processes, including intracellular transport.

## Materials and Methods

### Caenorhabditis elegans strains

The *C. elegans* strains used in this study are listed in the Table (Table S1). Some strains were provided by the CGC, which is funded by the NIH Office of Research Infrastructure Programs (P40 OD010440). Worms were maintained at 22°C. For the observation of cytoplasmic actin signal, the MG589 strain, expressing GFP::utrophin, was incubated at 25°C overnight before the observation to match the condition with the myosin knocked-down experiment, in which the worms were incubated at 25°C overnight.

### RNA interference

RNAi was conducted by directly injecting dsRNA into the worms^52^ for *nmy-2* and feeding for *perm-1*^76^. The dsRNA against *nmy-2* was synthesized using the primers with T7 and T3 promoters as follows: 5’-TAATACGACTCACTATAGGAAGTGCGTCATCTCTGGCTT-3’ and 5’-AATTAACCCTCACTAAAGGGAGGAACGACGAAGAACTCG-3’. The dsRNA synthesized from the genomic DNA of the N2 strain was dissolved into the TE buffer and stored at −20 °C until the injection. After the injection, the worms were incubated at 25°C for 16-20 h before the observations. For feeding RNAi of *perm-1*, the *Escherichia coli* clone for *C. elegans* PERM-1 T01H3.4 from the Ahringer library was used^77,78^. The bacterial culture (200 µL/plate) was spread onto 35 mm NGM agar plates containing 0.01 mM IPTG (isopropyl β-D-thiogalactopyranoside). Ten to twenty L4 worms were transferred onto each plate and incubated at 20 °C for 14-20 hours before observations.

### Imaging of *C. elegans* early embryos

To observe the fluorescent signal in the early embryos, we first dissected worms in 0.75× egg salt (118 mM NaCl, 40 mM KCl, 3.4 mM CaCl_2_, 3.4 mM MgCl_2_, 5 mM HEPES pH 7.2). The embryos from the dissected worms were mounted in 0.75× egg salt, which was placed on a 26 × 76-mm custom-made coverslip (Matsunami Glass). No coverslip was mounted on the embryos to eliminate the effects of deformation. The egg-mounted coverslips were set to the inverted microscope (Olympus, IX71), which was equipped with the CSU-X1 spinning-head (Yokogawa). The images were captured with a 60× silicon-immersion objective lens (Olympus, UPlanSApo, 60x/1.30Sil), a 2.0× intermediate magnification lens, and an EM-CCD camera (Andor, iXon), which was managed by the NIS elements software (Nikon). The exposure times were 300 ms for GFP::RAB-5 and mCherry::RAB-5, 180 ms for DHC-1::degron::GFP and GFP::utrophin, and 30 ms for 100 nm and 40 nm fluorescence beads. The experimental room was air-conditioned, and temperature was maintained at 21–23°C. The embryos which were confirmed to hatch were analyzed except for the myosin-perturbing experiments (*nmy-2* RNAi, NMY-2 injected, and blebbistatin treated).

### Tracking early endosome transport, artificial motor-cargo complex, and injected beads

Movements of early endosomes, artificial motor-cargo complex, and injected beads in 1-cell stage *C. elegans* embryo were analyzed by the custom-made software^79^ (Mark2). For the analysis of unidirectional movements, the tracked 2D trajectories were rotated to align the y-axis to the direction of the movements. The rotating angles were determined by the least squared fitting of individual 2D trajectories. The trajectories with the standard deviation in lateral position of less than 200 nm and more than 3-frame duration were categorized as unidirectional movements. The rotated 1D trajectories of unidirectional movements were fitted by a linear line, and the averaged speed was calculated. To calculate the diffusion coefficient of two-dimensional movements of beads, we fitted the MSD curves of individual trajectories by a linear function (*y = 4Dt, D* is the diffusion coefficient) using the time interval up to 1/4 of the whole tracking time. To calculate the MSD exponent of beads’ movements, we fitted the MSD curves of individual trajectories by the function y = *At^α^* (A is a fitting constant and α is the MSD exponent) using the time interval up to 1/4 of the whole tracking time. The mean diffusion coefficient and MSD exponent were calculated as the mean values of individual trajectories.

### Drug treatments

For the dynein degradation assay, the CA1215 strain expressing TIR1::mRuby and dhc-1::degron::GFP was used^53^. To degrade endogenous dynein, acetoxymethyl indole-3-acetic acid (IAA-AM), which is an analog of natural auxin IAA and can efficiently permeate the eggshell of *C. elegans*^55^, was used. Worms were dissected in the 0.75× egg salt, and the derived embryo was transferred into the egg salt containing 500 mM of IAA-AM.

To inhibit the activity of NMY-2 in the embryos, we increased the permeability of eggshells by knocking down the *perm-1* gene. In the observations of blebbistatin-treated embryos, the worms cultured on the *perm-1* feeding RNAi plate were dissected using 0.75×egg salt buffer supplemented with 2.5 µM blebbistatin (Sigma, B0560-1MG). After dissection, the worms were mounted using the same buffer.

### Optical flow analysis

To analyze fluctuating movements of actomyosin networks in the cytoplasm, we calculated the optical flow in the time-lapse confocal fluorescence images of GFP::utrophin in *C. elegans* embryo using the Farneback method^80^, which is implemented in the OpenCV library (www.opencv.org). The images of embryos from the MG589 worm strain (GFP::utrophin) were captured at 180 ms intervals. The captured images were rotated so that the embryo’s long axis appeared parallel to the horizontal axis. The images acquired with 180 ms intervals were averaged over four frames to calculate flows. After the averaging, the optical flow between the image at the n-th frame and at (*n*+4)-th frame was calculated. The mean flow speed was calculated by averaging the lengths of all the flow vectors. For the comparison among the myosin perturbed conditions, the averaged flow speed from 100 images from each embryo was used for each condition. Because the worms for *nmy-2* RNAi condition were incubated at 25℃ for 16-20 h as described in the “RNAi section.” Thus, the worms for other conditions were also incubated at 25℃.

### Coefficient of variation analysis of actin network

To analyze the spatial pattern of actin networks in the cytoplasm, we calculated the coefficient of variations of the signal of GFP::utrophin with varying the window size of *n* pixel (*n* = 2 - 25). After scanning the image with the window, the mean and standard deviation of intensity in the window and the following CV were calculated. After scanning the whole area of the image, the mean CV of the image was calculated. To obtain the CV of pixel-shuffled images, we randomly shuffled the intensity values in the pixels, and the CV was calculated for the shuffled image as described above.

### Plasmid construction and protein purification

The minimum motile construct of dynein (hsDHC-1_382k_), the heavy meromyosin (HMM) fragment of *C. elegans* NMY-2 (1-1364 aa), full-length MLC-4, and full-length MLC-5 were purified from mammalian cultured cells (FreeStyle 293-F cell, Invitrogen). For purifying hsDHC-1_382k_, the truncated 382 kDa heavy chain of human cytoplasmic dynein I was used as previously described^28^. For the purification of the NMY-2 complex, the plasmids coding NMY-2, MLC-4, and MLC-5 were co-transfected. The sequence for the amplification of the NMY-2 fragment was determined based on a previous study^59^. The sequences of the three genes were amplified from the cDNA of N2 strain worms. The amplified sequences were inserted into pcDNA3.4 vector for *nmy-2* and pcDNA5/FRT/TO for *mlc-4* and *mlc-5*, with the insertion of 6×His-tag for *nmy-2* and Streptavidin binding peptide (SBP)-tag for *mlc-4* and *mlc-5* in the upstream of each gene. The plasmids were transfected to the cultured cells with Polyethylenimine (PEI)^81^. The cells were harvested 72 h after the transfection, flash-frozen with liquid nitrogen, and stored at −80°C. For the purification, the frozen cells were thawed and suspended in lysis buffer (50 mM HEPES-KOH, 150 mM NaCl, 1 mM EGTA, 10% (w/v) sucrose, and pH 7.2) supplemented with the ProteoGuard EDTA-free protease inhibitor cocktail (Clontech, 635673). The suspended solution was put on ice and homogenized using a sonicator (Qsonica, Q125) with 60% amplitude and 1-s ON/1-s OFF pulses. The total sonicating time was 10 minutes. The homogenized solution was centrifuged at 75,000 rpm for 15 minutes (Beckman, TL100.3). The supernatant was loaded onto a StrepTactin Sepharose column (IBA) with a bed volume of 1 mL. After the loading, the column was washed with lysis buffer. The proteins were eluted with lysis buffer supplemented with 2.5 mM desthiobiotin (IBA, 2-1000-002) and flash frozen by liquid nitrogen. The concentration of purified proteins was determined by the Bradford method using TaKaRa Bradford Protein Assay Kit (Takara Bio, T9310A).

Tubulin used for *in vitro* observations was purified from porcine brain tissue through 2 successive cycles of polymerization and depolymerization. A high-molarity PIPES buffer was used to remove contaminating microtubule-associated proteins^82^. To prepare fluorescently labeled microtubules, we labeled tubulin with Cy-3 (PA23001, GE Healthcare) for the optical trap experiments or ATTO647N (AD 647N, ATTO TEC) for *in vitro* motility experiments. For *in vitro* motility experiments, the tubulin labeled with biotin (Tokyo Chemical Industry, S0956) was also used for immobilizing microtubules. Labeling of tubulins was performed according to the published method^83^. To prepare microtubules double-labeled with biotin and ATTO647N, non-labeled tubulins, ATTO647N-labeled tubulins, and biotinylated tubulins were mixed at the molar ratio of 20:1:1. Microtubules were polymerized in BRB80 (80 mM K-PIPES (pH 6.8), 1 mM MgCl_2_, and 1 mM EGTA) in the presence of 1 mM GTP and 5 µM taxol at 37°C for 15 minutes and then stabilized with additional taxol (50 µM for 15 minutes). Microtubules were then pelleted at 80,000 rpm in TLA-120.1 rotor at 25°C, and the pellet was resuspended in BRB80 with 50 µM taxol. The microtubules prepared were stored at room temperature and used within two weeks.

### Preparation of PEG-coated fluorescent beads

The fluorescent beads for analyzing cytoplasmic movements were coated by PEG to prevent non-specific binding of cytoplasmic proteins^84^. For the coating, the solution of carboxylated beads was diluted into 400 µL of 1× Phosphate-Buffered Saline (PBS) (Takara Bio, T900) containing 100 mg/mL PLL-g-PEG (SuSoS) with the particle number of 1.4 ×10^12^ particles for 40 nm bead (Invitrogen, F8794) and 7.2 ×10^11^ particles for 100 nm bead (Invitrogen, F8801). After the dilution, the mixtures were sonicated for 10 s and incubated for 1 h with 5 s sonication every 20 minutes. The beads were washed and concentrated using a centrifugal filter (Millipore, UFC510024), with 1× PBS. The particle density was adjusted to 0.7 ×10^10^ particles/µL for both beads.

### Preparation of BG-modified beads and conjugation with SNAP-fused motors

To prepare benzylguanine-modified beads, carboxylated 100 nm beads (Invitrogen, F8801) were subjected to modification using BG-NH_2_, as previously described^28^. A 10 nM fluorescent bead solution was mixed with 250 µM of sulfo-NHS (N-hydroxysulfosuccinimide) (Thermo Scientific, 22980) and 250 µM of EDC (1-ethyl-3-(3-dimethylaminopropyl)carbodiimide hydrochloride) (Thermo Scientific, 24510) in Borate buffer (100 mM Borate-NaOH, pH 8.5). The mixture was incubated for 30 minutes at room temperature. Following sonication, 650 µM of BG-NH_2_ (New England Biolabs, S9148) was added to the mixture, and the reaction proceeded for 2 hours at room temperature. To remove excess chemicals, the solution underwent three washes using a centrifugal filter (Millipore, UFC510024) with 50 mM Tris-HCl (pH 8.0). Finally, the particle density was adjusted to 1.44 ×10^10^ particles/µL by adding 50 mM Tris-HCl (pH 8.0). Beads less than 2 weeks from the modification were used for the conjugation with dynein. For the conjugation of dynein to BG-modified bead, the 120 nM hsDHC-1 and 1 nM BG-beads were mixed in 1×PBS and incubated at 25°C for 30 minutes. After the conjugation, the mixed solution was stored at 4 °C until injections.

### Gonad injection of recombinant proteins or motor-cargo complex

Purified proteins or the artificial motor-cargo complex were diluted using 1× PBS (Takara Bio, T900) and loaded into custom-made microneedles prepared with the P1000IVF micropipette puller (Sutter Instrument). In the case of NMY-2 injection, the concentration of total protein, including the HMM fragment of NMY-2 and two light chains (MLC-4 and MLC-5), was set to 100 µg/mL. Young adult worms were placed on a thin layer of 2% agarose (Lonza, SeaKem LE agarose) prepared on a 24 × 55-mm coverslip (Matsunami Glass, C024551, thickness No.1). After covering the worms with halocarbon oil, the coverslips were mounted on the inverted microscope (Carl Zeiss, Axiovert135). The protein solution supplemented with Texas Red-labeled dextran with a molecular weight of 3k (Invitrogen, D3329) was injected into the gonad of worms using the microinjector (Eppendorf, FemtoJet). After the completion of injections, 3 µL of M9 buffer^85^ was placed on the oil to release the worms. After the release, the worms were transferred to a new NGM plate and incubated at 22 °C for at least 3 h to incorporate the injected components into the embryos through oogenesis.

### Imaging and analysis of *in vitro* motility of the dynein-bead complex

*In vitro* motility assays were performed using an inverted microscope (Ti, Nikon) equipped with Epi-fluorescence imaging optics^86^. Image acquisition was performed using NIS-Elements software (ver 4.51.00, Nikon) with a 100× objective (1.49 NA, Nikon), fluorescence filters (chroma 49008 for microtubule; chroma ET705/72m for dynein beads), and EMCCD camera (Andor, iXon Ultra). Time-lapse images were acquired with 100 ms exposure without intervals. Flow chambers were assembled by attaching a glass coverslip (Fisher, 12548A) on a glass slide (Matsunami Glass, FF-001) using a pair of double-stick tape aligned in parallel with ∼3 mm separation. Before assays, the glass surface was pre-coated with a mixture of biotinylated PEG and non-biotinylated PEG (Laysan Bio Inc.) to minimize nonspecific protein binding^87^. The chamber was pre-coated sequentially with the following reagents: 1.0 mg/mL α-casein (Sigma-Aldrich, C6780) for 2 minutes, 0.2 mg/mL neutravidin (Thermo Fisher, A2666) for 5 minutes, and ATTO647N-labeled, biotinylated microtubules for 8 minutes. Then, dynein-conjugated, BG-coated beads which were diluted in Assay Buffer (12 mM K-PIPES pH 7.0, 1 mM MgCl_2_, 1 mM EGTA, 50 mM KCl, 2 mM DTT, 1 mM ATP, 20 µM taxol, and oxygen scavenging mix (4.5 mg/mL glucose, 0.20 mg/mL glucose oxidase and 0.035 mg/mL catalase)) supplemented with 0.25 mg/mL α-casein, was infused into the chamber.

### Instrumentation for optical trapping experiments

Optical trapping was performed essentially as described^88^. Beads were trapped and positioned over polarity-marked microtubules by a focused Nd:YAG laser (1064 nm, ATLAS, Coherent) using a 60×/NA1.45 oil immersion objective lens (Olympus, custom-made for maximum transmission of IR light; PlanApo). For rapidly moving the trap center, the trapping laser was passed through an electro-optic deflector (Quantum Technology, Model 41) and controlled with a multifunction generator (NF Corporation, WF1974). The bead was illuminated diagonally by a collimated red laser beam (644 nm; Excelsior-640C-60-CDRH, Spectra Physics). The light scattered by the bead was gathered by the same objective lens and passed through a custom-made perforated mirror (5 mm in diameter when viewed from the optical axis, Sigma Koki) placed below the objective lens^89^. The dark-field image of the bead was projected onto a quadrant photodiode (Hamamatsu Photonics, S994-13) coupled to a differential amplifier (Sentec, OP711G-2). The bead position was recorded at a sampling rate of 24 kHz and anti-aliasing filtered to 11 kHz before digitization by a 16-bit A/D module (National Instruments, USB-6251). Cy3-microtubules and dark-field images of the beads (just for manipulation) were simultaneously visualized by using TIRFM with a diode-pumped solid-state laser (532 nm; µGreen-SLM, JDS Uniphase). The images were split into two side-by-side images of Cy3-microtubule and beads using a dichroic mirror and projected onto a cooled EMCCD detector (DU-897E, Andor Technology). The exposure time was 70 ms. The sample was mounted on a piezoelectric stage (Physik Instrumente, P-517.3CL). Linearity between the monitored and actual displacement of the bead from the trap center was verified up to ±200 nm by applying a sinusoidal movement to the piezo stage and analyzing the response of the photodiode sensor. The bead position was calibrated for each bead by moving the photodiode sensor in a stepwise manner^90^. The trap stiffness was calculated for each bead based on the amplitude of the thermal diffusion using the equipartition theorem. The trap stiffness calculation was cross-checked by using power spectral measurements. A sinusoidal movement was applied to the sample to obtain a calibration peak, as previously described^91^.

### Preparation of dynein-coated beads for optical trap experiments

Twenty microliters of 1% carboxylate-modified 200 nm polystyrene beads (Invitrogen, F8811) were incubated with 0.6 µL of anti-SNAP antibody (New England Biolabs, P9310S) in a total of 50 µL BRB80 buffer (80 mM PIPES-KOH pH 6.8, 2 mM MgCl_2_, and 1 mM EGTA) for 20 minutes at room temperature. The bead surface was then blocked with 6–7 mg/mL casein for 10 minutes, followed by extensive washing with BRB80 buffer using a tabletop centrifuge (Eppendorf, 5424R). The collected beads were resuspended with 50 µL of BRB80 buffer and sonicated for 15 s (Velvo-Clear, VS-25). The resulting anti-SNAP beads were mixed with dynein for 15 minutes at room temperature and washed with BRB80 buffer. For single-molecule measurement, dynein was mixed with the bead at a sufficiently low concentration such that the binding fraction of the beads onto microtubules was less than 30%. This ensures that >90% of the beads that moved on a microtubule were driven by a single molecule^92^.

### Optical trapping experiments

A flow chamber was constructed using two coverslips (18 × 18 mm and 24 × 32 mm, thickness No.1; Matsunami Glass) and Parafilm (Heathrow Scientific LLC) to create a chamber that was 3 mm wide and 18 mm long. The Parafilm was used as a spacer and melted by placing the flow chamber on a 98°C hot plate. For immobilizing microtubules, the flow chamber was first coated with 10 µg/mL anti-β-tubulin antibody (Santa Cruz, SC-58884) in BRB80 buffer, allowed to adsorb for 5 minutes, and blocked with 1% (w/v) Pluronic F-127 (Sigma-Aldrich, P2443) in BRB80 buffer. After washing with 0.6–0.7 mg/mL casein in BRB80 buffer, the flow chamber was incubated with microtubules labeled by Cy3 in BRB80 buffer for 5 minutes. After washing with casein solution, the chamber was filled with imaging solution containing motor protein, 12 mM PIPES-KOH, pH 6.8, 2 mM MgSO_4_, 1 mM EGTA, 25 mM K-acetate, 10 µM paclitaxel, 0.7 mg/mL casein, 2 mM DTT, 25 mM glucose, 21.3 U/mL glucose oxidase, 800 U/mL catalase, and 1 mM ATP. The trapping experiments were performed at 24 ± 1°C. The force and duration were analyzed manually using custom-made LabVIEW (National Instruments)-based software. The average maximum force was determined by averaging the force corresponding to the maximum height of each peak. Data were low-pass filtered to 25 Hz to measure the maximum force prior to the analysis.

## Acknowledgements

Some strains were provided by the *Caenorhabditis* Genetics Center funded by the NIH Office of Research Infrastructure Programs (P40 OD010440). The SWG001 strain was kindly provided by Dr. Nathan Goehring (Francis Crick Institute). IAA-AM was kindly gifted by Dr. Takefumi Negishi (National Institute of Genetics). This work was supported by JSPS KAKENHI (grant numbers 18KK0202 to A.K. and T.T., 19K16094 and 24K09405 to T.T., and 18H02414 to A.K.).

## Author contributions

Conceptualization, T.T; Methodology, T.T., K.S., K.F., and A.K.; Investigation, T.T., K.S., and K.F; Writing – Original Draft, T.T.; Writing – Review & Editing, T.T., K.S., K.F., and A.K.; Funding Acquisition, T.T. and A.K.; Resources, A.K.

## Declaration of interests

The authors declare no competing interests.

## Supplemental information

Supplementary Document S1. Figures S1-S5 and Table S1

## Notes

### Competing Interest Statement

The authors have declared no competing interest.

