## Supplementary Document S1 for "Active fluctuations of cytoplasmic actomyosin networks facilitate dynein-driven intracellular transport along microtubules"

### Supplementary Figures

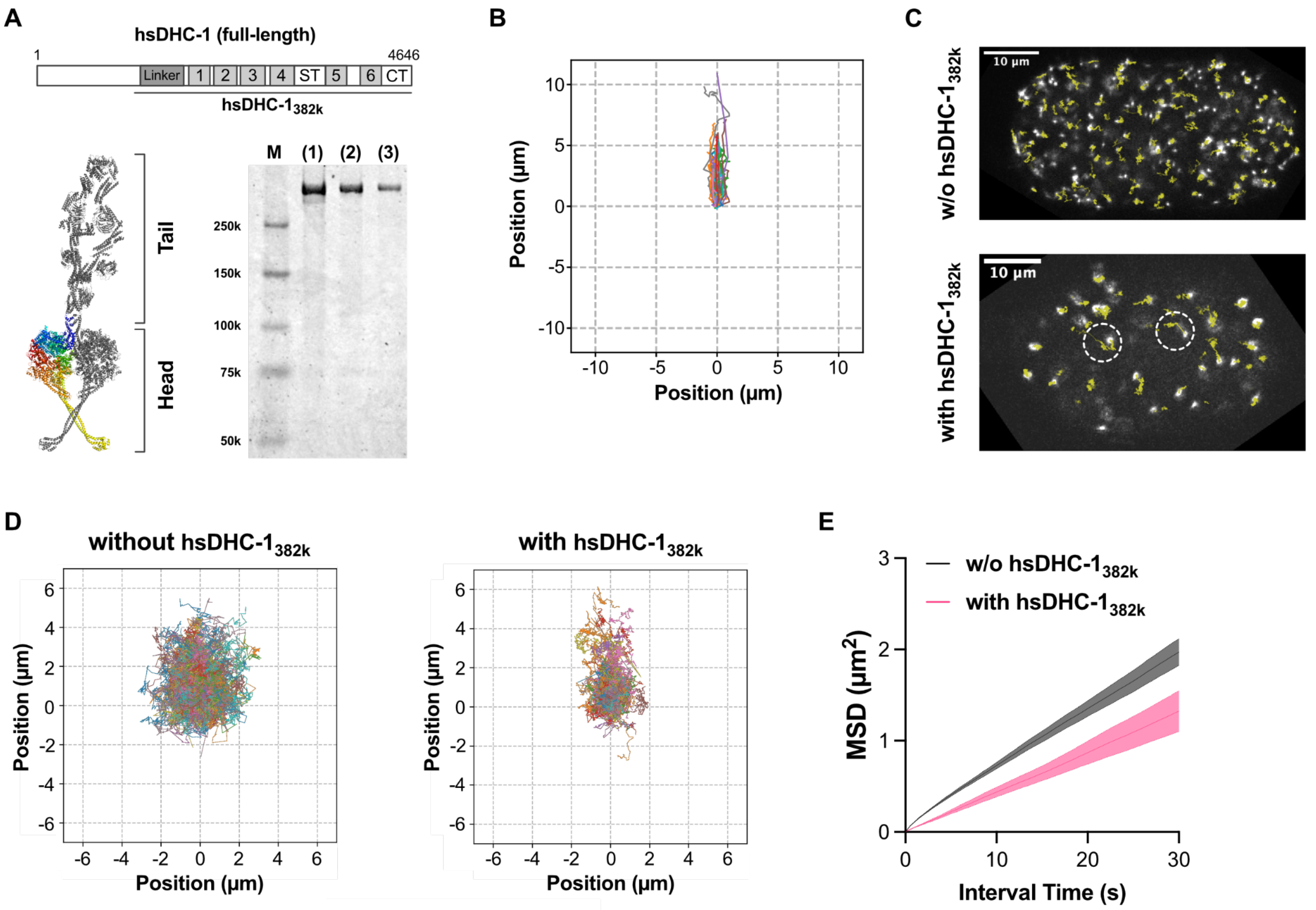


**Figure S1. Direct motility comparison between *in vitro* and *in vivo* using the artificial motor-cargo complex.** (A) A schematic diagram showing the truncated human dynein construct used in this study (top). The construct contains the linker and the whole motor domain. The numbers indicate AAA domains from AAA1 to AAA6. "ST" denotes the stalk domain. The protein structure presented in the bottom left shows a full-length dynein complex (PDB: 5NVU). The gel image shows SDS-PAGE of purified hsDHC-1$_{382k}$ (bottom right). The lane M denotes the marker. The following three lanes show the samples with different amounts of hsDHC-1$_{382k}$ (950 ng in (1), 475 ng in (2), and 230 ng in (3)). (B) Two-dimensional trajectories of BG-beads conjugated with hsDHC-1$_{382k}$, the motions of which are categorized as unidirectional. The trajectories are rotated so that the primal moving direction is aligned with the vertical axis. (C) Two-dimensional trajectories of BG-beads (yellow lines) in the embryos. The trajectories are superimposed on the raw images of BG-beads. The trajectories surrounded by the dashed-line circles indicate the unidirectional movements in the presence of hsDHC-1$_{382k}$. (D) Two-dimensional trajectories of benzylguanine(BG)-modified beads with or without dynein (hsDHC-1$_{382k}$) conjugation. The number of trajectories is 1,815 (without hsDHC-1$_{382k}$) and 852 (with hsDHC-1$_{382k}$), respectively. The trajectories were rotated so that the direction of the most prominent displacement, calculated from the least-square fitting with a linear line, was aligned with the positive direction of the y-axis. (E) MSD plots of BG-beads, obtained from the trajectories shown in (A).

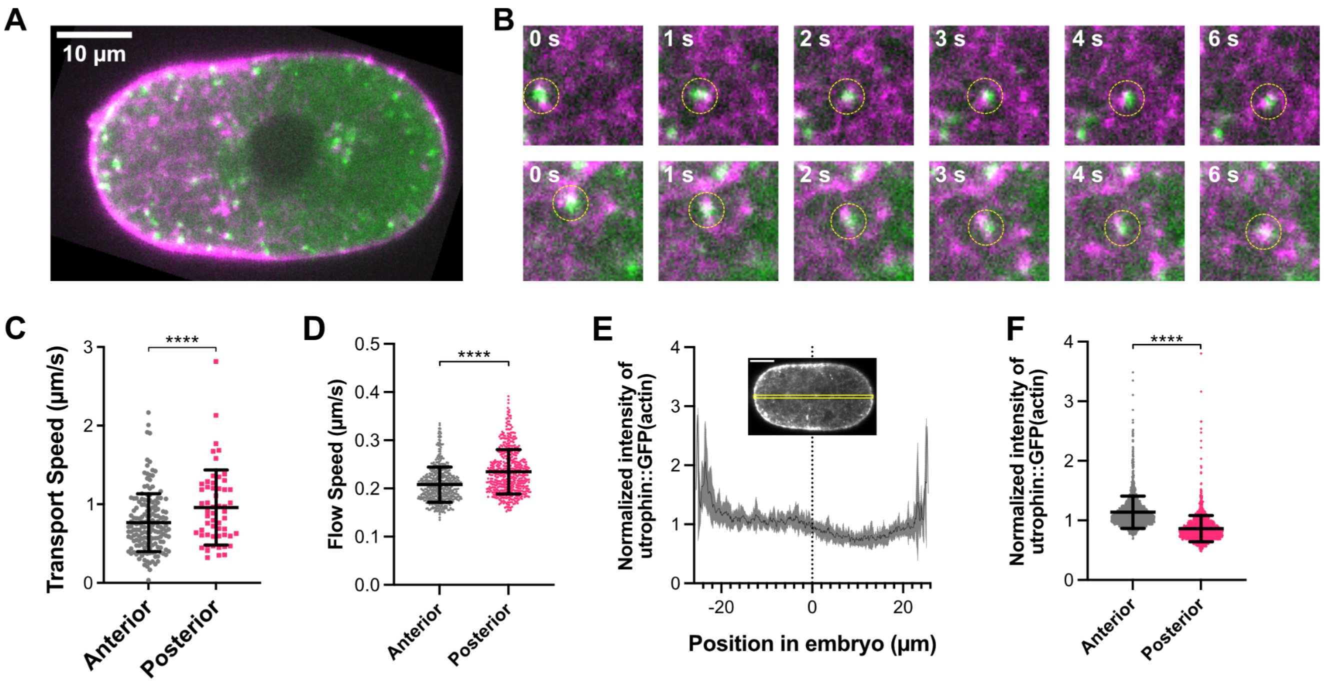


**Figure S2. Dynein-driven transport of early endosomes in *C. elegans* embryo.** (A) The image of a 1-cell stage embryo simultaneously expressing RAB-5::GFP (green) and LifeAct::mKate2 (magenta). (B) Examples of unidirectionally-moving early endosomes associating with actin networks in the cytoplasm. Time-series images of RAB-5 positive vesicles (green) associated with actin networks visualized by GFP::utrophin (magenta) are shown. The times elapsed from the starting time point are indicated in each image. The yellow dotted circles denote the RAB-5 positive vesicles. (C) The comparison of transport speed between the anterior and the posterior sides of embryos. Each plot represents the mean speed of individual trajectories. The *p*-value shown in the graph was obtained from Welch's *t*-test. (D) Mean optical flow speed calculated from the time-series images of GFP::utrophin. The flows in the anterior and the posterior sides were calculated separately. Each plot represents the mean flow speed at the single time points. The bars indicate the mean and the standard deviation. The statistical significance of the difference in mean was tested using Welch's *t*-test. (E) The intensity profile of GFP::utrophin (actin) along the anterior-posterior axis. A typical image example of the embryo is shown in the inset. The origin of the x-position is the geometrical center of the embryo, and the left side corresponds to the anterior side. The black line represents the intensity averaged over 10 images from 10 embryos. The gray area indicates the standard deviation. (F) The mean normalized intensity of the GFP::utrophin in the anterior and posterior sides. Each plot represents the intensity of individual pixels. The bars indicate the mean and the standard deviation.

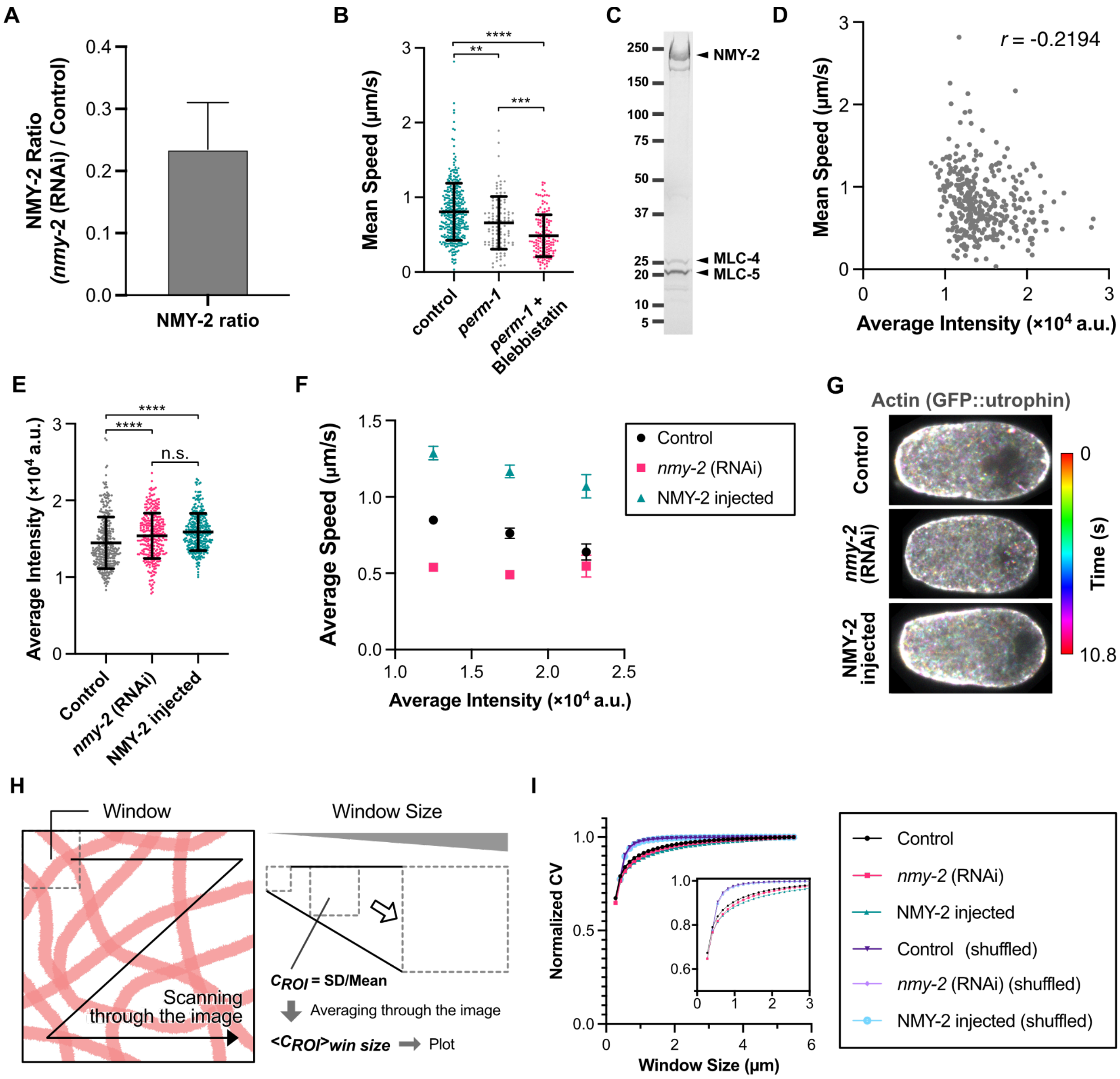


**Figure S3. Effects of modulating NMY-2 level in *C. elegans* early embryos.** (A) The ratio of endogenously-tagged GFP::NMY-2 in *nmy-2* (RNAi) embryos compared to the control embryos. For the calculation, the intensities of autofluorescence obtained from the WT (N2) strain were subtracted from the intensity values in the GFP channel for both embryos. The number of embryos analyzed was 12 for the control, 13 for *nmy-2* (RNAi), and 12 for N2, respectively. The bar indicates the bootstrap standard error (SE). (B) The comparison of transport speeds of early endosomes among the control, *perm-1* (RNAi), and *perm-1* (RNAi) + Blebbistatin embryos. Each point represents the mean speed calculated by the linear fitting to individual trajectories. The bars indicate mean and SD ($6.6 \pm 3.5 \times 10^2$ nm/s for *perm-1* (RNAi) and $4.9 \pm 2.8 \times 10^2$ nm/s for *perm-1* (RNAi) + Blebbistatin) The number of trajectories analyzed was 348 from 13 embryos (control, the same data shown in Figure 2), 132 from 4 embryos (*perm-1* (RNAi)), and 168 from 6 embryos (*perm-1* + Blebbistatin), respectively. The statistical significance was tested by the Kruskall-Wallis test followed by Dunn's Multiple

Comparison test (**: $p < 0.001$, ***: $p < 0.001$, and ****: $p < 0.0001$). (C) The gel image of SDS-PAGE of NMY-2 complexes (NMY-2, MLC-4, and MLC-5). The complex was purified from mammalian cells. (D) Relationship between the mean transport speed and the mean intensity of unidirectionally moving early endosomes in the control embryos. The original data of mean speeds was the same as that shown in Figure 2. (E) Comparison of mean intensities of early endosomes obtained from the control, *nmy-2* (RNAi), and NMY-2 injected embryos. The original data used were the same as those shown in Figure 2 and Figure 3. The bars indicate the mean and SD. The statistical significance was tested by the Kruskall-Wallis test followed by Dunn's Multiple Comparison test (n.s.: not significant, ***: $p < 0.001$, and ****: $p < 0.0001$). (F) Comparison of the mean transport speed of early endosomes, where the data were divided into three groups based on the mean intensity values. (G) Temporal-color-coded images of actin (utrophin::GFP) in the unperturbed (control), *nmy-2* (RNAi), and NMY-2 injected embryos. The time is indicated by the color, as shown in the right bar. (H) Schematic diagram showing the coefficient of variations (CV) analysis of actin intensity. The standard deviation of the intensity divided by the mean intensity was calculated with the defined window size. (I) Comparison of the window size dependency of CV analysis of the control, *nmy-2* (RNAi), and NMY-2 injected embryos. The data from the images with pixel shuffling are presented, too. The inset indicates the curves on a shorter length scale. The images used for the calculation were 500 from 10 embryos for all conditions. The CVs are normalized by the maximum values of each condition.

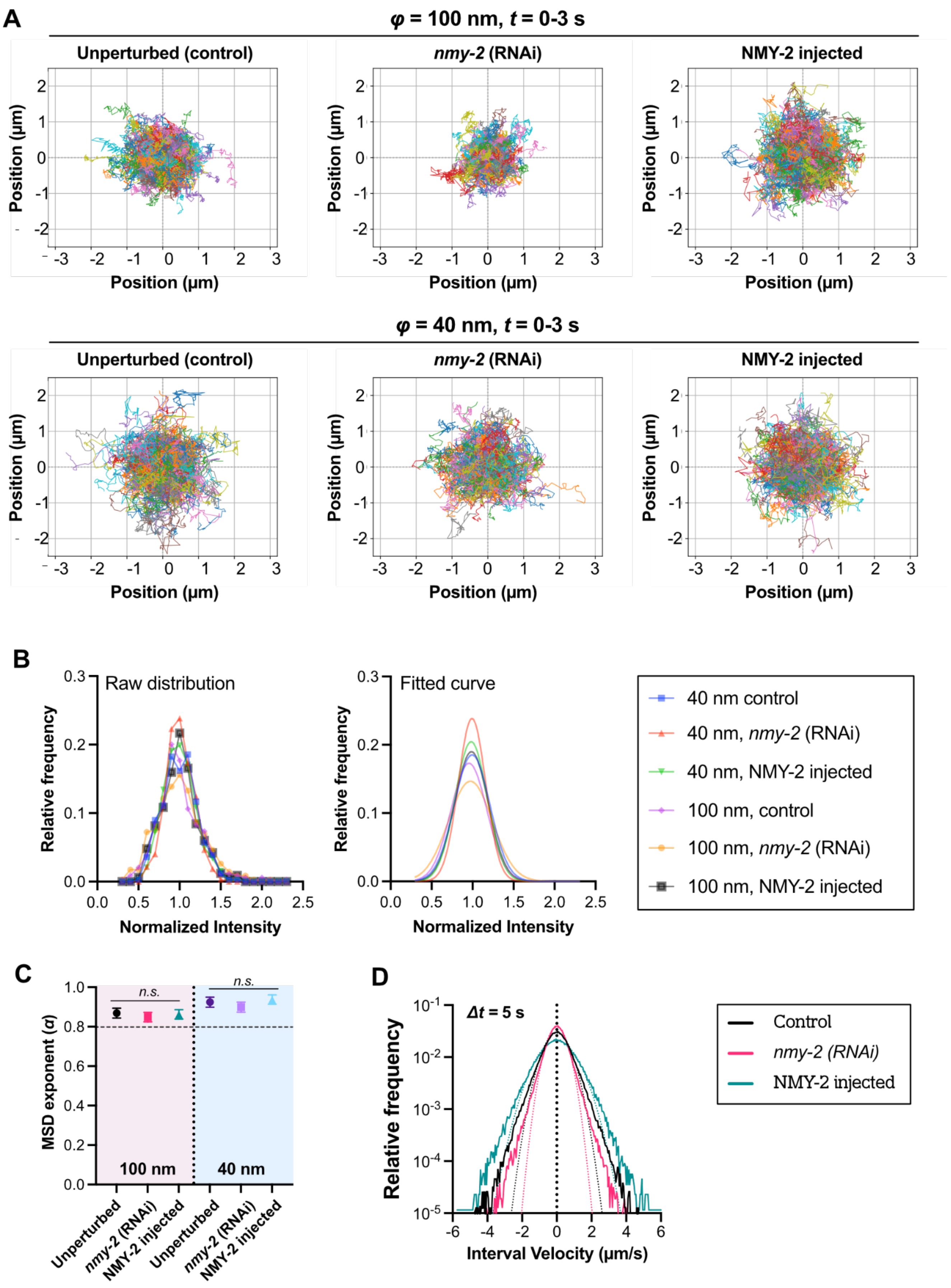


**Figure S4. Movements of beads incorporated into the *C. elegans* embryos.** (A) Two-dimensional trajectories of beads incorporated into the 1-cell stage embryo of *C. elegans* over 3 seconds. The origin of the position is the starting point of each trajectory. (B) Intensity distribution of beads used for MSD calculation. The normalized raw data (left) and the fitted curve by a single Gaussian (right) are shown. (C) Comparison of the MSD exponent calculated from the log-log plots of MSD shown in Figure 4C. Mean and SEM are shown. The statistical significance among the conditions was checked by the Kruskal-Wallis test. (D) Distributions of interval velocity ($\Delta t$ = 150 ms) from the unperturbed (control), *nmy-2* (RNAi), and NMY-2 injected. The solid lines show the raw distributions. The dashed lines indicate the fitting by a single Gaussian.

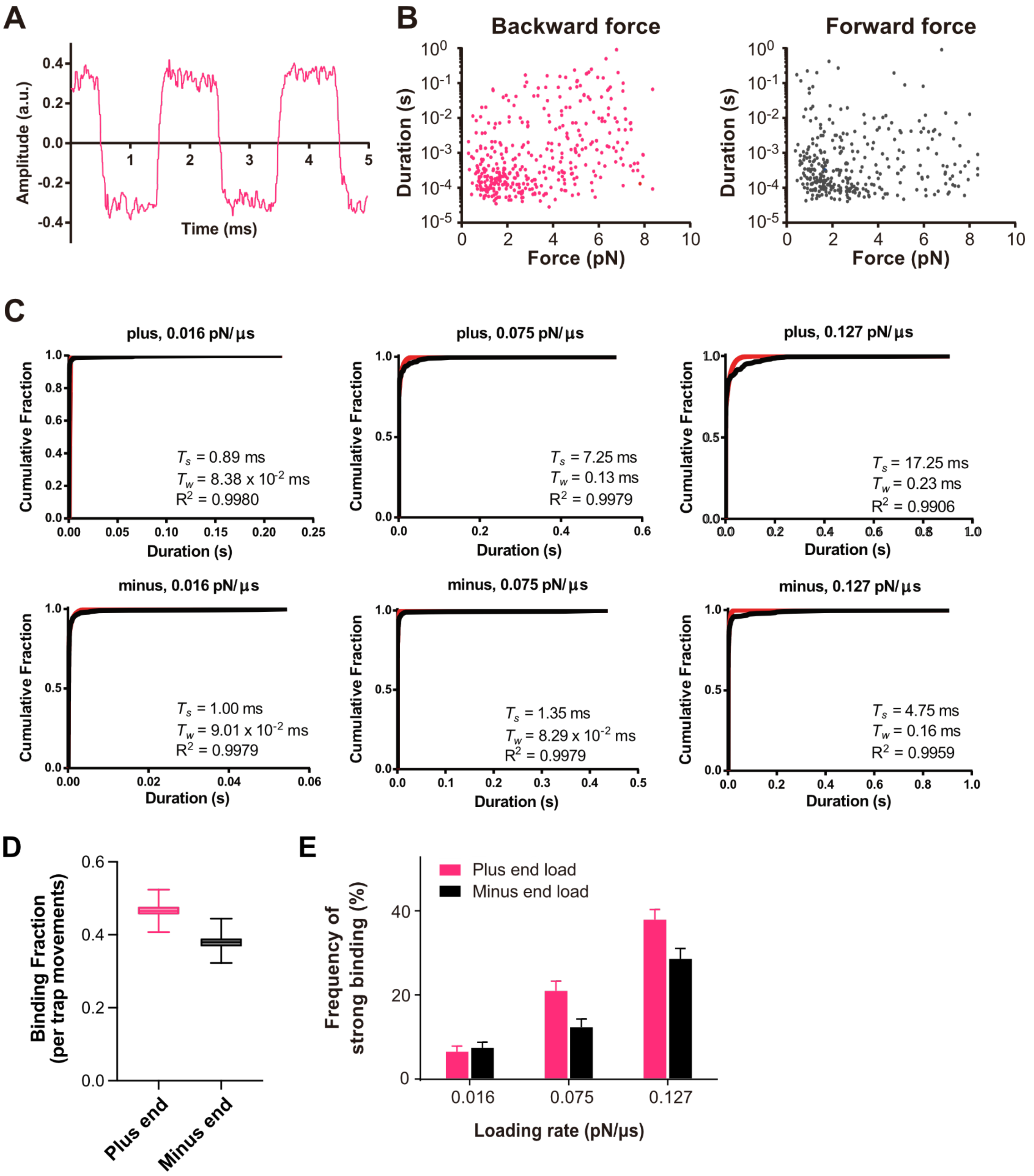


**Figure S5. Force response of dynein measured by the optical trap.** (A) Representative time series of the displacement of a trapped bead, recorded by an oscilloscope. The bead image was projected onto a quadrant photodiode, and the displacement was measured while the trap center was rapidly moved using the electro-optic deflector. The maximum scanning speed was measured as 240 nm per ~70 µs (a loading rate of 0.127 pN/µs at a trap stiffness of 0.035 pN/nm). (B) Raw data of the duration versus force. The data were acquired at a loading rate of 0.127 pN/µs. (C) Cumulative plots of the duration of hsDHC-$1_{382k}$ on microtubules at different loading rates (black lines) and the associated fits by a sum of two exponential functions (red lines) (n = 287–392 per plot). The fit obtained from the sum of two exponential functions yielded two different time constants corresponding to short ($T_2$) and long ($T_1$) binding events, and the values are shown at the lower right of each graph. For long binding events, the time constant for backward movements of the trap center was increased from 0.9 to 17.6 ms (20-fold), whereas that for forward scan was increased from 1.0 to 4.8 ms (4.8-fold). (D) Frequency of observing binding events during the scanning movements of the trap center. The total number of trap center movements was 840. The bars indicate the bootstrap SEM. (E) Frequency dependency of strong bindings on loading rates. The frequency is defined as the ratio of the strong bindings to all binding events.

**Table S1.** The *C. elegans* strains used in this study

| | Strain | Genotype | Comment | Figure | Source | Reference |
|---|---|---|---|---|---|---|
| **1** | N2 | *C. elegans* wild isolate | | Figure 1<br>Figure S1<br>Figure 3<br>Figure S3 | CGC | Brenner, 1974[1] |
| **2** | RT122 | *unc-119*(ed3) III; *pwIs20* [*pie-1p*::*gfp*::*rab-5* + *unc-119*(+)] | | Figure 2 | CGC | Sato et al., 2005[2] |
| **3** | RT1043 | *unc-119*(ed3) III; *pwIs403* [*pie-1p*::mCherry::*rab-5* + *unc-119*(+)] | | - | CGC | Sato et al., 2008[3] |
| **4** | SWG001 | *gesIs001*[*Pmex-5*::Lifeact::mKate::*nmy-2*UTR, *unc-119*(+)] | | - | | Reymann et al., 2016[4] |
| **5** | CA1215 | *dhc-1*(*ie28*[*dhc-1*::degron::*gfp*]) I; *ieSi38*[*sun-1p*::TIR1::mRuby::*sun-1* 3'UTR + Cbr-*unc-119*(+)] IV | | Figure 1<br>Figure S1 | CGC | Zhang et al., 2015[5] |
| **6** | CAL2621 | *unc-119*(ed3) III; *pwIs20*[*pie-1p*::*gfp*::*rab-5*]; *unc-119*(ed3) III; *gesIs001*[*Pmex*-5::Lifeact::mKate::*nmy-2*UTR, *unc-119*(+)] | RT122 × SWG001 | Figure S2 | - | This study |
| **7** | CAL2661 | *unc-119*(ed3) III; pwIs403 [*pie-1p*::mCherry::*rab-5* + *unc-119*(+)];*dhc-1*(ie28[*dhc-1*::degron::*gfp*]) I; *ieSi38*[*sun-1p*::TIR1::mRuby::*sun-1* 3'UTR + Cbr-*unc-119*(+)] IV | CA1215 × RT1043 | Figure S2 | - | This study |
| **8** | LP162 | *nmy-2*(*cp13*[*nmy-2*::*gfp* + LoxP]) I. | | Figure S3 | CGC | |
| **9** | MG589 | *xsSi3* [gfp::*utrophin* + Cbr-*unc-119*(+)] | | Figure 3<br>Figure S3 | CGC | Tse et al., 2012[6] |
